# Intrinsically ball retrieving wolf puppies reveal standing ancestral variation for human-directed play behaviour

**DOI:** 10.1101/744862

**Authors:** Christina Hansen Wheat, Hans Temrin

## Abstract

Domestication dramatically alters phenotypes. Standing variation among ancestral populations often drives phenotypic change during domestication, but some changes are caused by novel mutations. Dogs (*Canis familiaris*) engage in interspecific play with humans and it has specifically been suggested that the ability to interpret social-communicative behaviour expressed by humans is a novel dog-specific skill. Thus, wolves (*Canis lupus*) are not expected to engage in interspecific play with a human based on social-communicative cues. Here we report the observation of three eight week-old wolf puppies spontaneously responding to social-communicative behaviours from a stranger by retrieving a ball. This unexpected and novel observation has significant implications for our understanding and expectations of the genetic foundations of dog behaviour. Importantly, our observations indicate that behavioural responses to human social-communicative cues are not unique to dogs. This suggests that, while probably rare, standing variation in the expression of human-directed behaviour in ancestral populations could have been an important target for early selective pressures exerted during dog domestication.

## Introduction

Domesticated animals express dramatic phenotypic alterations compared to their ancestral species [1,2]. While phenotypic change can be attributed to novel mutations, a growing body of evidence suggest that evolutionary change relies heavily upon standing genetic variation [3,4]. Indeed, though few novel mutations with large effects account for some phenotypic differences between domestic and ancestral populations [4,5], animal domestication was likely initiated by selection on standing genetic variation within ancestral populations [4]. The potential for domestic phenotypes to derive from existing variation has been well demonstrated in the farm fox project [6,7], where strong selection regimes based on observed variation in the behavioural trait tameness (i.e. reduced aggression and increased docility) among pre-selection foxes brought about rapid occurrence of classic morphological phenotypes associated with domestication. Knowing whether the basis for traits selected upon during early domestication are variants from ancestral populations or unique mutations arising during domestication is central to developing our understanding of the domestication process. For instance, wild species expressing variation for the trait tameness are arguably more likely to be successfully domesticated compared to species that do not [8]. Therefore, disentangling whether phenotypic change in domesticates is caused by novel mutations or selection on standing ancestral variation is important if we are to advance our understanding of the domestication process and its generalities across species.

The dog (*Canis familiaris*), which was domesticated from the grey wolf (*Canis lupus*) at least 15,000 years ago [2], show extreme phenotypic variation as a species. Present day dogs are bred for highly breed-specific requirements for behaviour and morphology [9,10], and while a significant amount of the resulting variation is believed to originate from standing genetic variation in ancestral populations [11], novel mutations have had a significant impact during breed formation [4]. For instance, black coat colour [12,13], chondrodysplasia (foreshortened limbs [5]) and brachycephaly (pathologically short muzzle [14]) are traits that have occurred in modern dogs through novel mutations. An additional example comes from a genome wide analysis of genetic difference between dogs and wolves, which identified dogs as having an increased copy number of the amylase locus (*AMY2B*), which was argued to be a novel adaptation to a starch-rich diet in early-domesticated dogs [15]. However, investigation of a wider range of individuals revealed standing variation in amylase copy numbers in wolves, thereby shifting the *AMY2B* example from being a novel mutation important in domestication, to yet another example of selection upon standing variation as an essential substrate for domestication [16]. This critical distinction has important implications for hypothesizing how domestication could have taken place. Thus, the *AMY2B* example illustrates the importance of including enough observations to detect existing variation among wolves and thereby avoiding over-interpreting the uniqueness of traits expressed in dogs.

While much progress has been made in studying the morphological and physiological differences between wolves and dogs, understanding the basis and origins of behavioural variation have proven more elusive [17]. One behavioural skill that has been suggested to be novel in dogs is interspecific social competence [18–20]. Specifically, it has been posited that, unlike wolves, dogs possess unique skills to interpret human cues [18] and that these skills might have arisen after the domestication process from the grey wolf had been initiated [20,21]. The ability to interpret human social cues has received considerable interest from researchers comparing behaviour in dogs and wolves. However, due to substantial differences in testing procedures, environmental factors and interpretation of results, consensus among these studies is lacking [22–26]. Consequently, whether wolves have the ability to interpret human social cues, or whether this is a novel dog-trait, remains unresolved.

Here we focus upon human-directed play behaviour, which has been reported in some domesticated species [27,28], including dogs [29–32]. Dogs can interpret human play cues and adjust their behavioural repertoires when playing with a human instead of a conspecific [31,33]. Within a domestication context, wherein animals have been selected for greater tolerance of and interactions with humans, interspecific human-directed play behaviour represents a highly relevant behaviour to address. However, to date only one study exists comparing human-directed playfulness in a domesticated species and its ancestral proxy species [34], and studies on human-directed play behaviour in wolves have never been attempted.

Here we report on the spontaneous expression of human-directed play behaviour, in the form of ball retrieving for a stranger, in eight week old, hand-raised wolves. Our observations occurred during a test in which puppies, with no prior training, are vocally encouraged to retrieve a ball and thus respond to social-communicative behaviours from a human they had never met before. Based on the existing literature, we expected that human-directed play behaviour is a novel trait that occurred during the domestication of dogs and that wolves therefore would not respond to interspecific social-communicative behaviours or engage in human-directed play with a stranger.

## Materials and methods

### a) Study animals

From 2014 to 2016 we hand-raised three litters of European grey wolves (N = 13) at Tovetorp Zoological Research Station, Stockholm University, Sweden. The wolf litters from 2014, three females and two males, and 2015, two males, were full siblings. The 2016 wolf litter consisted of four males and two females and was not related to the wolf litters from 2014 and 2015. Hand-raising was initiated from the age of 10 days, before eye opening, for all litters. By choosing a hand-raising set-up, we were able to minimize environmental bias, including maternal effects, which is well-documented to affect the development of behavioural patterns [35–37]. Wolves were raised within litters and extensively socialized, which included 24-hour presence of human caregivers for the first eight weeks. All wolves were reared under standardized conditions across all three years. Hand-rearing was initiated in identical indoor rooms and at the age of five weeks the wolves were given access to smaller roofed outdoor enclosures. After a habituation period of one week, the wolves were given additional access to a larger fenced grass enclosure at six weeks of age. Thereafter the wolves had free access to the indoor room and the two enclosures during the day and access to the indoor room and the roofed enclosure during the night. Behavioural observations began at 10 days of age and behavioural testing was initiated at 6 weeks of age. Hand-raising, testing procedures and exposure to the new environments were standardized over all three years, which included the implementation of rules to assure that rearing was standardized across all caregivers. This included that wolves were never disciplined or trained. Wolves never met strangers until their vaccination program was completed at eight weeks of age and, importantly, not until the completion of the test in which the observations of this study were recorded. Behavioural testing prior to eight weeks of age did not include other people than the caregivers.

### b) Behavioural sampling

Wolves were tested in the Puppy Mental Assessment (PMA) at eight weeks of age. The PMA is a standardized behavioural test battery developed by the Swedish Working Dog Association based on the need to offer dog breeders a standardized test to describe puppy behaviour in specific situations. The results from the PMA can serve as a tool for dog breeders to choose suitable new owners for their puppies. As such, puppies are tested before they leave the breeder at seven to nine weeks of age. The PMA consists of 42 standardized tests situations covering behaviours in four main groups: 1) Social play with a stranger, here the puppy assessor, 2) Object play and object interest 3) Social comfortableness and fearfulness and 4) Interest in strangers, here the puppy assessor, including greeting. The puppy is tested in a novel room or an enclosure. The PMA starts with the owner or familiar person (in this study CHW or HT) placing the puppy in the middle of the test room, in which the puppy assessor is already present (but neutral), and then leaves the room swiftly. The whole test takes approximately 10-15 minutes. The subtest in which our observations occurred is related to social play with a stranger. In this test the puppy assessor throws a tennis ball across the room and calls the puppy back, encouraging it to retrieve the ball. Retrieving and cooperation is measured by the puppy’s willingness to return the ball to the puppy assessor and is scored on a 1 to 5 scale (Table 1). The test is repeated three consecutive times.

**Table 1.** Behavioural scoring. Cooperation in the three consecutive retrieving tests is measured on a scale from 1 to 5, where 1 is no cooperation and 5 is full cooperation.

| *Score* | *Behaviour* |
| --- | --- |
| 1 | The puppy shows no interest in the ball |
| 2 | The puppy plays with the ball on its own, but aborts |
| 3 | The puppy plays with the ball on its own, but ignores the puppy assessor’s call |
| 4 | The puppy responds to the puppy assessor’s call, initiates retrieving but releases the ball |
| 5 | The puppy responds to the puppy assessor’s call and retrieves the ball to her |

### Ethical statement

Daily care and all experiments were performed in accordance with relevant guidelines and regulations under national Swedish Law. The experimental protocols in this study were approved by the Ethical Committee in Uppsala, Sweden (approval number: C72/14). Facilities and daily care routines were approved by the Swedish National Board of Agriculture (approval number: 5.2.18-12309/13). All wolves were born in animal parks and CITES certified with at least F2 status.

## Results

Three wolves, all from the 2016 litter, fully retrieved the ball at least two times and one of those wolves fully retrieved the ball all three times (Score: 5, Fig 1, Supplemental Video 1). On one occasion one of the wolves fully retrieving the ball two times also played with the ball, but ignored the puppy assessors call (score: 3, Supplemental Video 2). One wolf from the 2014 litter and one from the 2016 litter showed some interest in playing with the ball on at least one trial but aborted (Score: 2). Eight wolves (four from the 2014 litter, both from the 2015 litter and two from the 2016 litter) showed no interest in the ball on any of the three trials (Score: 1, Supplemental Video 3).

**Fig. 1.**
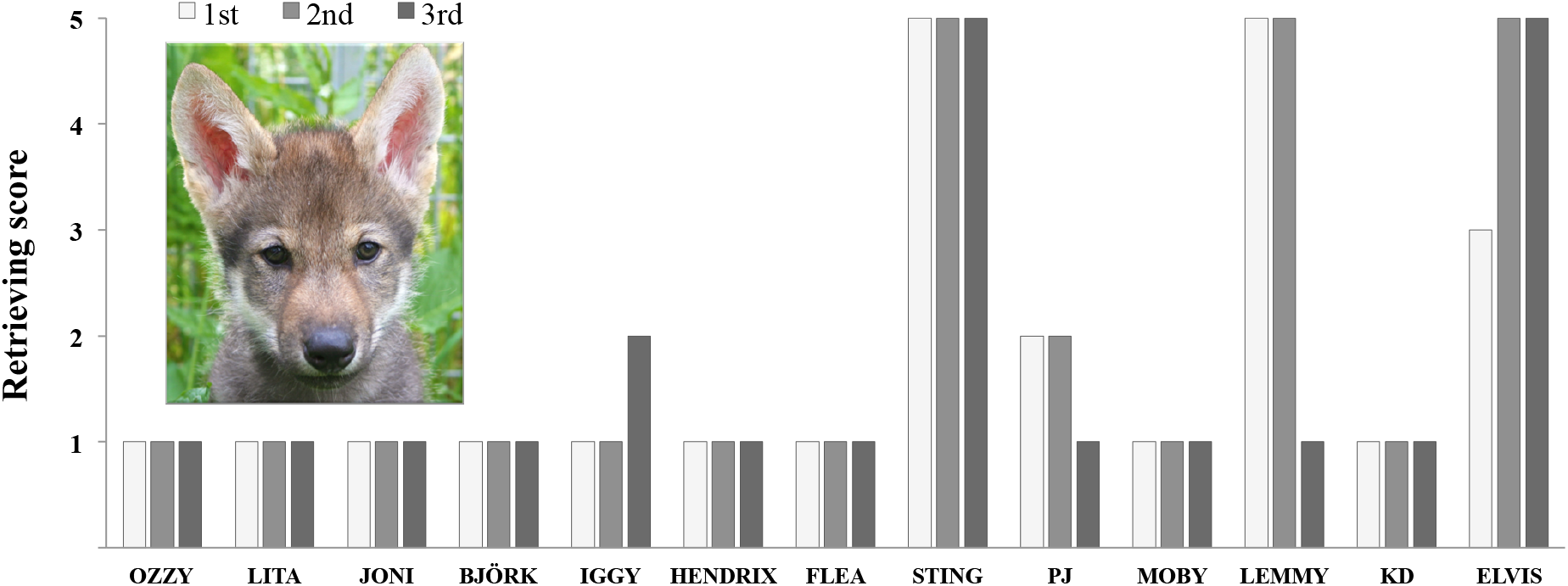
Behavioural scores. Cooperation scores in the retrieving test 13 wolves on three consecutive trials (shading from light to dark with the 1st trial being light, 2nd medium and 3rd dark). Behaviour is scores on a scale from 1-5. Only scores 4 and 5 include partial or full retrieving, respectively. Photo credit: Christina Hansen Wheat

## Discussion

Here we provide the first empirical evidence that a behaviour thought unique to dogs, namely interspecific play with a human based on social-communicative cues, actually exists in wolves. Our finding is surprising given that the ability to interpret social-communicative behaviour expressed by humans has been suggested to be a novel dog trait [18–21]. Importantly, our results suggest that, while probably rare, standing variation in the expression of human-directed behaviour, including play, in ancestral populations could have been an important target for early selective pressures exerted during dog domestication.

Our novel observations of three wolf puppies retrieving a ball are highly relevant for the on-going discussion on the effect domestication has on behaviour and further have significant implications for our understanding and expectations about the genetic foundations of the behaviours in modern day dogs. Specifically, in relation to current attempts to reveal the genomic basis of behavioural changes during domestication [16,38,39], our observations indicate signatures of selection for human-directed behaviour in dogs are likely to be weak and prone to false positives [*sensu lato* 40,41]. This is because 1) we must now consider that selection likely acted upon standing variation in interspecific social-communicative behaviour in wolves, 2) this behaviour almost certainly has a polygenic genetic architecture, and 3) samples sizes in recent genomic studies are small and therefore lacking sufficient power to detect the resulting expected selection dynamics.

In sum, we argue that in order to answer questions about the evolutionary foundation of dog behaviour, research attention should refocus away from solely conducting direct species comparisons, and include studies upon whether or not specific behavioural variation inherently exists among wolves. Identifying such instances has important ramifications upon expectations of how dog domestication proceeded.

## Acknowledgments

We wish to thank Stockholm University for funding this study, our hand-raisers Patricia Berner, Anna Björk, Marjut Pokela, Linn Larsson, Charles Gent, Åsa Lycke, Erika Grasser, Joanna Schinner, Yrsa Andersson, Christoffer Sernert and Mija Jansson, the staff at Tovetorp Zoological Research Station, the puppy assessors Ingrid Tapper and Inga-Lill Larsson of the Working Dog Association for their help conducting the tests, and Wouter van der Bijl and Christopher W. Wheat for helpful comments on the manuscript.

